# Identification of TEX101 functional interactome through proteomic measurement of human spermatozoa homozygous for the missense variant rs35033974

**DOI:** 10.1101/315739

**Authors:** Christina Schiza, Dimitrios Korbakis, Keith Jarvi, Eleftherios P. Diamandis, Andrei P. Drabovich

**Author notes:** Correspondence should be addressed to: A. P. Drabovich, Ph.D., Department of Laboratory Medicine and Pathobiology, University of Toronto, 60 Murray Street; Room L6-201-4, Toronto, ON, M5T 3L9, Canada., or, E. P. Diamandis, Ph.D., M.D., Mount Sinai Hospital, 60 Murray St [Box 32]; Flr 6 - Rm L6-201-1, Toronto, ON, M5T 3L9, Canada.

## Abstract

TEX101 is a testis-specific cell-surface protein expressed exclusively in the male germ cells and a validated biomarker of male infertility. Mouse TEX101 was found essential for male fertility, and was suggested to function as a cell surface chaperone involved in maturation of proteins required for sperm migration and sperm-oocyte interaction. However, the precise functional role of human TEX101 is not known and cannot be studied *in vitro* due to the lack of human germ cell lines. Here, we genotyped 386 healthy fertile men and sub-fertile patients for a common and potentially deleterious missense variant rs35033974 of *TEX101*, and identified 52 heterozygous and 4 homozygous patients. We then discovered by targeted proteomics that the variant allele rs35033974 was associated with near-complete degradation (>97%) of the corresponding G99V TEX101 form, and suggested that spermatozoa of homozygous patients could serve as a knockdown model to study TEX101 function in humans. Differential proteomic profiling with label-free quantification measured 8,046 proteins in spermatozoa of eight men and identified 8 cell-surface and 9 secreted testis-specific proteins significantly down-regulated in four patients homozygous for rs35033974. Substantially reduced levels of testis-specific cell-surface proteins potentially involved in sperm migration and sperm-oocyte fusion (including LY6K and ADAM29) were confirmed by targeted proteomics and western blotting assays. Since recent population-scale genomic data revealed homozygous fathers with biological children, rs35033974 is not a single pathogenic factor of male infertility in humans. However, median TEX101 levels in seminal plasma were found 5-fold lower (*P*=0.0005) in heterozygous than in wild-type men of European ancestry. We conclude that spermatozoa of rs35033974 homozygous men have substantially reduced levels of TEX101 and could be used as a model to elucidate the precise TEX101 function, which will advance biology of human reproduction.

**Non-standard abbreviations:**

TEX101: Testis-expressed sequence 101 protein
LY6K: Lymphocyte antigen 6 complex locus K
ADAM29: A disintegrin and metalloproteinase domain-containing protein 29
DPEP3: Dipeptidase 3
BH-adjusted t-test: Benjamini-Hochberg-adjusted t-test
FDR: False discovery rate
FWHM: Full width at half maximum
GPI: Glycosylphosphatidylinositol
LC-MS/MS: liquid chromatography - tandem mass spectrometry
LFQ: Label-free quantification
MS: Mass spectrometry
mAb: Monoclonal antibody
MWU: Mann Whitney Unpaired t-test
PRM: Parallel reaction monitoring
ROC AUC: Receiver operating characteristic area under the curve
SCX: strong cation exchange chromatography
SP: seminal plasma
SNV: Single nucleotide variation
SRM: Selected reaction monitoring
WT: wild-type

## INTRODUCTION

Recent-omics studies identified 1,079 human genes with exclusive expression in testis (1). While function of many of testis-specific proteins is not known, it may be assumed that these proteins have unique and specialized roles in spermatogenesis and fertilization. Mutations, natural knockouts or deleterious single nucleotide variations in testis-specific genes could lead to severe male infertility with spermatogenesis arrest, low sperm concentration, abnormal spermatozoa morphology or low motility. In addition, mutations in testis-specific genes involved in sperm transit, acrosome reaction and sperm-oocyte interaction may result in no apparent phenotypical issues and male infertility which cannot be explained with routine diagnostics (2-4).

We previously discovered and validated a testis-specific protein TEX101 as a seminal plasma biomarker for the differential diagnosis of azoospermia and male infertility (5-9). The functional role of TEX101 is not known, but based on mouse models it was suggested as a spermatozoa cell-surface chaperone involved in the maturation of four cell-surface proteins from the ADAM family (10-12). *Tex101* knockout in mice resulted in absolute male sterility but normal sperm concentration, morphology and other phenotypical characteristics (10, 11). In the absence of TEX101 protein, ADAM 3-6 proteins were not properly processed and degraded. However, mouse data could not be translated into human studies since ADAM3, ADAM5 and ADAM6 genes are non-coding pseudogenes, while ADAM4 is not present in the human genome (13). Due to the lack of stable human male germ cell lines, TEX101-dependent proteins could not be identified in humans.

As an alternative, we suggested that the functional role of TEX101 could be studied in human clinical samples, such as spermatozoa. Our previous work on TEX101 levels in seminal plasma revealed a small population of men with high sperm count, but unexplained infertility and very low levels of TEX101 protein in seminal plasma and spermatozoa (8). In this work, we hypothesized that some genomic alterations, such as natural knockouts or deleterious single nucleotide variations, could result in undetectable or low levels of TEX101 protein. We suggested that spermatozoa obtained from such patients could be used as knockout or knockdown models to identify proteins degraded in the absence of TEX101 and discover the functional interactome of TEX101 in humans. Unlike immunoprecipitation approaches to reveal direct and strong physical interactions, profiling of the whole proteome of spermatozoa lacking TEX101 could identify the whole functional interactome including weak and transients interactions. Collectively, such data could support in humans the previously suggested function of TEX101 as a cell-surface chaperone (10).

## EXPERIMENTAL PROCEDURES

### Study design and statistical rationale

The objectives of this study were to identify potential genomic alterations which could impact levels of TEX101 protein, and verify those levels experimentally in human spermatozoa samples. We suggested that differential proteomic profiling of spermatozoa with such genomic alterations could identify proteins degraded in the absence of TEX101 and thus discover the functional interactome of TEX101 in humans. According to power calculations, differential profiling of spermatozoa of 4 wild-type and 4 rs35033974 homozygous men could identify proteins down-regulated at least 2.4-fold, assuming 80% power, α = 0.05, 1.8% coefficient of variation for log2-transfromed LFQ intensity values and a one-tailed t-test. Power calculations were done with G^*^Power software (version 3.1.7, Heinrich Heine University Dusseldorf). GraphPad Prism (v5.03) was used to generate scatter plots, perform statistical analysis, and calculate the area under the Receiver Operating Characteristic curve (ROC AUC). Non-parametric Mann – Whitney U test was used to compare TEX101 levels in seminal plasma of wild-type and heterozygous men, and *P*-values <0.05 were considered statistically significant.

### Study population and sample collection

Semen samples (*N*=386) were collected with informed consent from patients with the approval of the institutional review boards of Mount Sinai Hospital (approval #08-117-E) and University Health Network (#09-0830-AE). Samples were obtained from healthy fertile men before vasectomy and individuals diagnosed with oligospermia or unexplained infertility. Clinical parameters are summarized in **Table 1**. The unexplained infertility group included men who were not able to father a pregnancy after one year of regular unprotected intercourse, with normal sperm concentration of greater than 15 million/mL. After liquefaction, semen samples were centrifuged three times at 13,000 g for 15 min at room temperature. Spermatozoa and SP were separated and stored at -80°C. Samples were analyzed retrospectively. For the differential proteomic analysis, spermatozoa samples were obtained from four men homozygous for *TEX101 c.296G*>*T* variant (rs35033974^*hh*^), diagnosed with oligospermia (*N*=2) and unexplained infertility (*N*=2), median age of 29.5 years, sperm concentration 2 – 30 million/mL, and TEX101 concentration in SP 3.5 – 47 ng/mL. Spermatozoa obtained from the age-matched wild-type (WT) fertile men referred for vasectomy (*N*=4, sperm concentration >15 million/mL and TEX101 concentration in SP of 8,000 – 12,500 ng/mL) were selected as a control group.

**Table 1.**
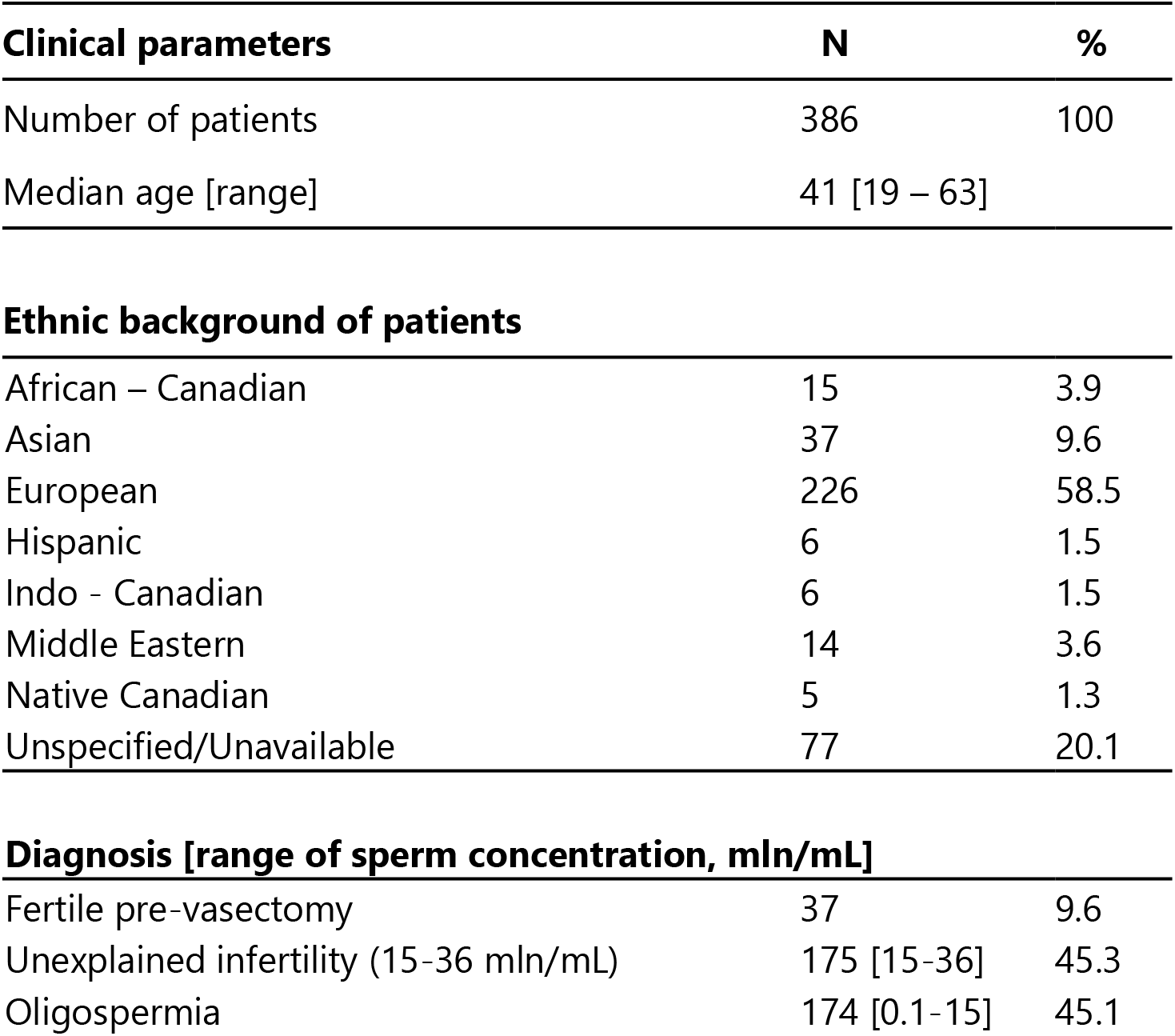
Clinicopathological variables of 386 patients.

### Extraction of genomic DNA from spermatozoa and *TEX101* genotyping

Genomic DNA was extracted from spermatozoa using QIAamp DNA Mini Kit (Qiagen Inc.). Spermatozoa were washed twice with phosphate-buffered saline (PBS). Cells were lysed in the presence of proteinase K and DNA bound to the membrane was washed and eluted. DNA purity and concentration were measured by spectrophotometer (NanoDrop 8000, Thermo Scientific). Forward (5’-ACAGGACTGAGACAGCCAT-3’) and reverse (5’-TCCAGGGTACCTGTGGTCTC-3’) primers were designed to amplify a 197 base pair fragment of *TEX101* gene encompassing the rs35033974 polymorphism. PCR was performed with 50 ng of genomic DNA, 1.2 units of Phusion High-Fidelity DNA polymerase (Thermo Scientific) in Phusion HF Buffer, 200 μM deoxynucleoside triphosphates and 0.5 μM primers using mastercycler thermal cycler (Eppendorf). PCR included an initial denaturation step at 98°C for 1 min, followed by 40 cycles of denaturation at 98°C for 10 s, annealing at 64°C for 30 s and extension at 72°C for 30 s, with a final extension at 72°C for 7 min. PCR products were confirmed with 1.5% agarose gel electrophoresis and purified with QIAquick PCR Purification Kit (Qiagen). Sequencing of PCR products (N=386 patients) was performed by the Centre for Applied Genomic (Hospital for Sick Children, Toronto).

### Sample preparation and protein digestion by endopeptidase Glu-C

Spermatozoa pellets were washed twice with PBS, lysed with 0.1% RapiGest SF (Waters, Milford, MA) in 50 mM ammonium bicarbonate and sonicated three times for 30 s. Cell lysates were then centrifuged at 15,000 g for 15 min at 4°C. Total protein in each spermatozoa or SP sample was measured by the bicinchoninic acid assay. Ten μg of total protein per patient sample in 50 mM ammonium bicarbonate were used for protein digestion. RapiGest SF 0.05% with 5 mM dithiothreitol at 65°C for 30 min were used to denature proteins (purified recombinant human rhTEX101 and proteins from spermatozoa and SP) and reduce disulfide bonds. Free thiols were then alkylated with 10 mM iodoacetamide in the dark for 40 min at room temperature. Protein digestion was completed overnight at 37°C in the presence of sequencing grade Glu-C enzyme obtained from Promega (1:20 Glu-C: total protein) and supplemented with 5% acetonitrile to enhance Glu-C activity. Digestion in the presence of ammonium bicarbonate at pH 7.8 ensured specific cleavage after glutamine residues. Trifluoroacetic acid (1%) was then used to inactivate Glu-C and cleave RapiGest SF detergent. Synthetic peptides representing the wild-type (AITIVQHSSPP<u>**G**</u>LIV*TSYSNYCE) and the G99V variant (AITIVQHSSPP<u>**V**</u>LIV*TSYSNYCE) forms of TEX101 were labeled with 13C5-, 15N-valine at the residue 102 and were used as internal standards spiked-in after digestion at final concentrations of 100 fmol/μL and 250 fmol/μL, respectively. Digests were desalted, and peptides were extracted by C18 OMIX tips (Varian Inc., Lake Forest, CA). Peptides were eluted into 3 μL of 70% acetonitrile with 0.1% formic acid and analyzed by an EASY-nLC 1000 nanoLC coupled to Q Exactive^TM^ Plus Hybrid Quadrupole-Orbitrap^TM^ Mass Spectrometer (Thermo Fischer Scientific).

### Development of Parallel Reaction Monitoring (PRM) assay for WT and G99V variant forms of TEX101 protein

To evaluate Glu-C specificity and efficiency of digestion, rhTEX101 protein was digested and analyzed in the data-dependent discovery mode, which confirmed specific generation of AITIVQHSSPP<u>G</u>LIVTSYSNYCE (m/z=1268.6) and AITIVQHSSPP<u>V</u>LIVTSYSNYCE (m/z=1289.6) peptides. Uniqueness of these peptides in the human proteome was confirmed by Basic Local Alignment Search Tool (http:/blast.ncbi.nlm.nih.gov/Blast.cgi). Following that, rhTEX101 and seminal plasma were digested with Glu-C and analyzed in the unscheduled targeted PRM mode. In the final optimized PRM method, heavy isotope-labeled peptide internal standards and an additional endogenous TEX101 peptide TAILATKGCIPE (m/z=637.3) were monitored (**Supplemental Table S1**). A four-step 16-min gradient was used: 20% to 40% of buffer B for 8 min, 40% to 65% for 2 min, 65% to 100% for 2 min, and 100% for 4 min. PRM settings were the following: 3.0 eV in-source CID, 17,500 MS2 resolving power at 200 m/z, 3×10^6^ AGC target, 100 ms injection time, 2.0 m/z isolation window, optimized collision energy at 27 and 100 ms scan times.

### Immunocapture-PRM measurements of WT and G99V TEX101

Total TEX101 protein was enriched from SP and spermatozoa using an in-house anti-TEX101 mouse monoclonal antibody 34ED556. Briefly, protein G purified 34ED556 monoclonal antibody was immobilized on NHS-activated Sepharose 4 Fast Flow beads (GE Healthcare). Fifty μL of beads (~25 μg of 34ED556) in 0.1% BSA were incubated overnight at 4°C with seminal plasma or spermatozoa lysate. After binding, beads were washed three times with tris buffer saline (50 mM Tris, 150 mM NaCl, pH 7.5) followed by washing with 50 mM ammonium bicarbonate. Proteins were digested overnight on beads using Glu-C. Supernatants were acidified with 1% TFA. Heavy peptides (200 fmoles of WT and 500 fmoles of G99V) were spiked into each sample after digestion. Digests were desalted, and peptides were measured by PRM assay. Raw files were analyzed with Skyline software (v3.6.0.10493), and the relative abundances of WT or G99V variant TEX101 forms were calculated using the light-to-heavy peptide ratios.

### Sample preparation for the differential proteomic analysis

Spermatozoa pellets from eight men (four WT and four rs35033974^*hh*^) were lysed with 0.1% RapiGest SF in 50 mM ammonium bicarbonate. Cell lysates were centrifuged at 15,000 g for 15 min at 4°C to remove debris, and total protein concentration was measured using BCA assay. Proteins (225 μg per sample) were denatured, reduced with 5 mM dithiothreitol, alkylated with 10 mM iodoacetamide and digested overnight with trypsin (Sigma-Aldrich) at 37°C.

### Strong-cation exchange chromatography fractionation

Off-line SCX fractionation was used to facilitate deep proteome analysis. Tryptic peptides were diluted with mobile phase A (0.26 M formic acid in 10% CAN at pH 2-3) and were loaded onto PolySULFOETHYL A^™^ column (2.1 mm ID×200 mm, 5 μm, 200 Å, The Nest Group Inc., MA). Peptides were separated with a 60-min three step HPLC gradient (Agilent 1100) and eluted at 200 μL/min with 1 M ammonium formate (0-15% for 5-25 min, 25% at 35 min and 100% at 50 min). Twenty-seven 400 μL fractions were initially collected, but then pooled into 13 fractions based on absorbance profiles.

### Protein identification by liquid chromatography - tandem mass spectrometry (LC-MS/MS)

Peptides of each SCX fraction were concentrated with C18 OMIX tips and analyzed by an EASY-nLC 1000 system coupled to a Q Exactive^TM^ Plus mass spectrometer in technical duplicates for each fraction (14, 15). Peptides were separated with a 15 cm C18 analytical column using a 90-min LC gradient at 300 nL/min flow rate. Full MS1 scans (400 to 1500 m/z) were acquired with the Orbitrap analyzer at 70,000 FWHM resolution in the data-dependent mode, followed by 12 data-dependent MS2 scans at 17,500 FWHM. Only +2 and +3 charge states were subjected to MS2 fragmentation.

### Data analysis and label-free quantification

XCalibur software (v. 2.0.6; Thermo Fisher Scientific) was utilized to generate raw files. For protein identification and label-free quantification, raw files were analyzed with MaxQuant software (version 1.5.2.8). MaxQuant searches were performed against the non-redundant Human UniprotKB/Swiss-Prot database (HUMAN5640_sProt-072016) at the false discovery rate (FDR) of 1.0%. Search parameters included: trypsin enzyme specificity, two missed cleavages, minimum peptide length of 7 amino acids, minimum identification of one razor peptide, fixed modification of cysteines by carbamidomethylation and variable modification of methionine oxidation and N-terminal protein acetylation. The mass tolerance was set to 20 ppm for precursor ions and 0.5 Da for fragment ions with top 12 MS/MS peaks per 100 Da. MaxLFQ algorithm facilitated label-free relative quantification of proteins (16). ‘ProteinGroups.txt’ file was uploaded to Perseus software (version 1.5.5.3) to facilitate statistical analysis (17). Proteins classified as ‘Only identified by site’, ‘Reverse’ and ‘Contaminants’ were filtered out, and LFQ intensities were log2-transformed. Missing LFQ values were imputed with the down shift of 1.8 and distribution width of 0.45 to ensure normal distribution, and average LFQ intensities for two technical replicates were calculated. A two-sample t-test with Benjamini-Hochberg false discovery rate-adjusted p-values was applied, and 5.0% FDR with calculated constant for variance correction s0=0.4 were used to select proteins differentially expressed in rs35033974^*hh*^ men. Data was visualized with the volcano plot. Significant up- or down-regulated proteins were filtered for the cell-surface and secreted proteins with the testicular tissue-elevated (tissue-enriched, group enriched and tissue-enhanced) expression according to the Human Protein Atlas, version 13 (1).

### Experimental design and rationale for development of Selected Reaction Monitoring (SRM) assays and quantification of candidate proteins

To quantify candidate proteins, we developed and applied Tier 2 SRM assays, as previously described (18-24). Briefly, LC-MS/MS peptide identification data was used to select proteotypic tryptic peptides and develop SRM assays. Choice of peptides was confirmed with the SRM Atlas (www.srmatlas.org). For each protein, peptides with 7-20 amino acids and without missed cleavages were chosen, and heavy isotope-labeled peptide internal standards were synthesized. Several unscheduled 30-min SRM methods were prepared and run with a pool of spermatozoa digest with TSQ Quantiva^TM^ triple quadrupole mass spectrometer (Thermo Scientific). The three most intense transitions were selected for each heavy or light peptide. Finally, 20 heavy and light peptides were scheduled within 2 min intervals during a 30-min gradient in a single multiplex SRM assay (**Supplemental Table S2**). The parameters for SRM assay included: positive polarity, 150 V declustering and 10 V entrance potentials, 300^°^C ion transfer tube temperature, optimized collision energy values, 20 ms scan time, 0.4 Q1 and 0.7 Q3 FWHM resolutions and 1.5 mTorr Q2 argon pressure. Since one rs35033974 homozygote spermatozoa sample was fully consumed in the discovery experiment, candidate proteins were quantified in three rs35033974 homozygote and four WT spermatozoa samples. Spermatozoa lysates (10 μg protein) were digested by trypsin. TEX101 and DPEP3 internal standards with trypsin-cleavable tags (500 fmoles of AGTETAILATK*-JPTtag and SWSEEELQGVLR*-JPTtag, respectively) were added before trypsin digestion, while 8 heavy isotope-labeled peptides without JPT tags were spiked after digestion (500 fmoles each). Light and heavy peptides were monitored with a scheduled 30-min multiplex SRM assay. Each spermatozoa sample was analyzed. Light-to-heavy ratio was used to calculate the accurate relative abundance of each candidate protein.

### Western blotting

Protein levels of TEX101, LY6K, ADAM29, and DPEP3 were assessed by Western blot analysis. Twenty μg of total protein from one rs35033974 homozygous and one WT spermatozoa lysate were loaded onto an SDS-PAGE gel (4% - 15%, Bio-Rad), and transferred onto PVDF membranes (Bio-Rad). After blocking, membranes were incubated overnight at 4°C with rabbit polyclonal antibodies against TEX101 and DPEP3 (HPA041915 and HPA058607, Atlas Antibodies), sheep polyclonal antibody against LY6K (AF6648, R&D Systems), mouse monoclonal antibody against ADAM29 (H00011086-M09, Abnova Corporation) and GAPDH antibody (AM4300, Thermo Fisher Scientific). The membranes were washed then incubated with goat anti-rabbit, donkey anti-sheep, and goat anti-mouse secondary antibody conjugated to horseradish peroxidase (Jackson ImmunoResearch). Proteins were detected with chemiluminescence substrate (GE Healthcare Life Sciences).

### Data availability

Raw mass spectrometry shotgun data and MaxQuant output files were deposited to the ProteomeXchange Consortium via PRIDE (www.ebi.ac.uk/pride/archive/login) with the dataset identifier PXD008333 and the following username: and password: IIdaf83v. PRM and SRM raw data were deposited to the Peptide Atlas with the dataset identifier PASS01112 (http://www.peptideatlas.org/PASS/PASS01112) and the following username: PASS01112 and password: NI5437g. Alternative link is ftp://PASS01112:NI5437g@ftp.peptideatlas.org. Processed Skyline files can be downloaded at Panorama Public (https://panoramaweb.org/3jbthK.url, password: jv7Z4#GC).

## RESULTS

### Database mining for loss-of-function variants of TEX101 gene

The Exome Aggregation Consortium (http://exac.broadinstitute.org), Genome Aggregation (http://gnomad.broadinstitute.org) and 1000 Genomes Project (www.internationalgenome.org) databases (25, 26) were examined for the presence of potential loss-of-function variants of the human *TEX101* gene. GnomAD database included genomic variants identified in 138,632 individuals of diverse ethnic background and revealed 166 potential loss-of-function variants of *TEX101* (**Supplemental Table S3**). Protein knockout or truncating variants, such as start loss, stop-gain and frameshift variants, were very rare (minor allele frequencies <0.003%), while some missense variants leading to single amino acid substitutions were much more frequent. One of such missense variants was rs35033974 (allele frequency 8.4%). Interestingly, rs35033974 was predicted as ‘deleterious’ by Polyphen (27), SIFT (28) and CADD (29) algorithms. The allele frequency of rs35033974 varied among different populations. It was more common in European non-Finnish population (12.4%), less common in Latino (5.2%), Ashkenazi Jewish (5.8%), South Asian (3.3%) and African (2%), and very rare in East Asian population (<0.00001%). Rs35033974 (c. 296 G>T) was localized within exon 4 and resulted in substitution of glycine to valine at position 99 (**Figure 1A**). Alignment of TEX101 protein sequences suggested that glycine-99 was a conserved residue in 17 of 19 mammals and thus could be intolerant to substitutions (**Supplemental Figure S1**). The high frequency of rs35033974 warranted its identification in our male infertility biobank. Spermatozoa samples obtained from heterozygous (22% genotype frequency in European population) and homozygous (1.6% genotype frequency) men were investigated.

**Figure 1.**
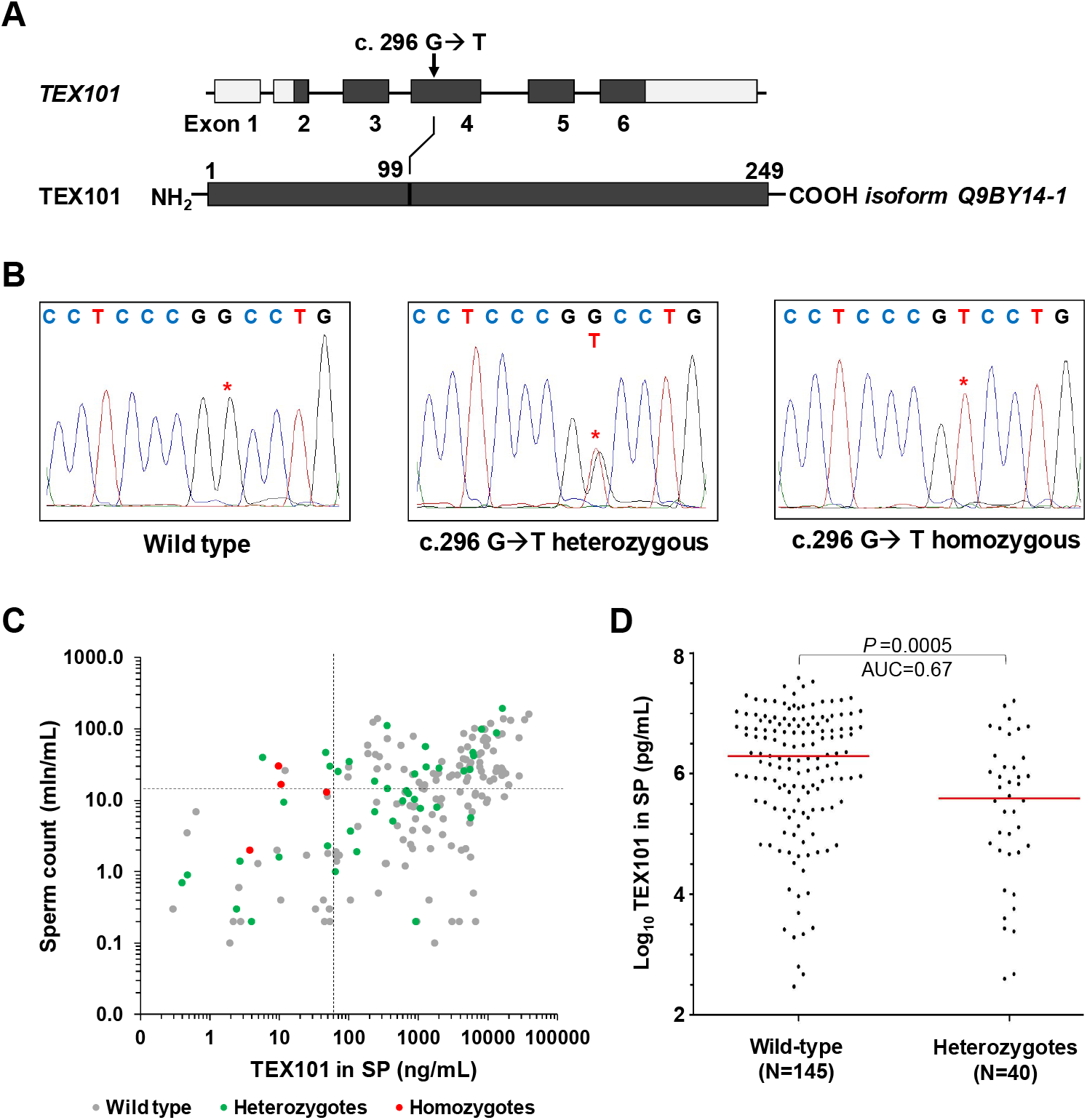
Identification of *rs35033974* variant in *TEX101* gene. (**A**) Schematic representation of TEX101 gene and protein sequence, showing the missense variant c.296 G>T, and the amino acid substitution p.99 G>V. (**B**) TEX101 DNA sequencing of WT *TEX101* patient (left panel), rs35033974 heterozygous patient (middle panel) and rs35033974^*hh*^ homozygous patient (right panel). (**C**) Scatter plot of sperm concentration and TEX101 levels in seminal plasma of 189 sub-fertile men (diagnosed with unexplained infertility or oligospermia) of European origin. WT, rs35033974 heterozygous and rs35033974^*hh*^ men are plotted in grey, green and red, respectively. Sperm concentration (15 million/mL) and TEX101 concentration (65 ng/mL) cut-off values are indicated by dotted lines in black. Variant rs35033974 allele frequency is higher for the groups with TEX101<65 ng/mL (58.3% for upper left group and 21% for lower left group), and lower for the groups with TEX101≥65 ng/mL (9.8% for lower right group, and 8.7% for upper right group). (**D**) Seminal plasma levels of TEX101 normalized by sperm concentration were significantly lower (fold change 4.0, MWU P=0.0005) for G99V heterozygous European men (median 390 ng/mL, *N*=40), as compared to WT men (median 1,949 ng/mL, *N*=145). Median values for each group are represented as horizontal lines.

### Identification of men heterozygous and homozygous for the rs35033974 variant

Genotypes for rs35033974 variant (c. 296 G>T) were determined in 386 men by amplification of spermatozoa DNA and sequencing analysis (**Table 1** and **Figure 1B**). Four heterozygous individuals (GT, genotype frequency 11%) were identified in the group of pre-vasectomy fertile men (*N*=37). In the group of patients with unexplained male infertility (*N*=175), 23 heterozygous (13%) and 2 (1.1%) homozygous patients (TT) were found. In the group of patients diagnosed with oligospermia (*N*=174), we identified 25 heterozygous (14%) and 2 homozygous (1.1%) patients. Investigation of patients with European ancestry (178 WT, 44 heterozygous and 4 homozygous men) revealed minor allele frequency of 11.5% and was similar to gnomAD frequency of 12.4%. In our cohort of European men, the minor allele frequency was higher for patients with unexplained infertility and oligospermia (12.7%) than infertile men pre-vasectomy (5.4%), but was not significant (Fisher’s exact test *P*=0.15).

### Impact of rs35033974 on TEX101 protein concentration in seminal plasma

We previously measured by ELISA concentration of total TEX101 protein in seminal plasma of 805 men (8). Cross-checking revealed that concentration of total TEX101 in seminal plasma of four rs35033974^*hh*^ homozygous patients was extremely low (3.5 to 47 ng/mL), despite of their medium-to-high sperm concentration (2 to 30 mln/mL; **Figure 1C**). Here, we also examined possible associations between rs35033974 heterozygous status and TEX101 levels in seminal plasma or sperm concentration in semen. No significant difference for sperm concentration was found for WT versus heterozygous European population (MWU *P*-value =0.94). However, levels of TEX101 in seminal plasma were significantly lower for rs35033974 heterozygous men (median 390 ng/mL, MWU P=0.0005, *N*=40), as compared to WT men of European population (median 1,949 ng/mL, N=145; **Figure 1D**). Since TEX101 concentration in seminal plasma may correlate with the number of spermatozoa in semen, we also investigated normalized TEX101 concentration. Thus, seminal plasma levels of TEX101 normalized by sperm concentration were significantly lower (4-fold change, MWU P<0.0001) for heterozygous men (median 40,738 attograms/cell, *N*=40), as compared to WT men (median 164,085 attograms/cell, *N*=145).

### Rs35033974 results in degradation of G99V TEX101 protein

To monitor the WT and G99V variant forms of TEX101 protein in spermatozoa lysate and SP, we opted to develop a targeted mass spectrometry assay (19, 30, 31). Since TEX101 digestion by trypsin generated a 38-amino acid peptide not suitable for bottom-up proteomic measurements, we explored alternative proteases, such as endopeptidase Glu-C (cleaves peptide sequences after aspartic and glutamic acids) and neutrophil elastase (cleaves after valine and alanine). Endopeptidase Glu-C was found the most suitable enzyme. Following optimization of Glu-C digestion protocol, we developed a PRM assay to monitor WT and G99V TEX101 peptides, as well as an additional endogenous “control” peptide which represented total TEX101. Sensitivity of PRM assay, however, was not sufficient to measure low levels of TEX101 in seminal plasma (<1.5 μg/mL).

To improve assay sensitivity, we developed an immuno-PRM assay based on the immunoenrichment of TEX101 by our in-house anti-TEX101 mouse monoclonal antibody 34ED556 coupled to sepharose beads (**Figure 2A**). Using immuno-PRM assay, we were able to identify barely detectable levels of G99V variant protein in one homozygous patient, while the other two homozygous patients had undetectable levels of G99V protein. We also selected one heterozygous patient with a very high concentration of TEX101 in seminal plasma (16.1 μg/mL), and were able to measure levels of both WT and G99V variant forms in his spermatozoa (Figure 2B). Interestingly, the abundance of a G99V variant form was substantially lower, as compared to the WT form. Using heavy-to-light ratios of both peptides, we estimated that the abundance of the G99V form was ~97% lower than predicted, assuming equal expression of both alleles.

**Figure 2.**
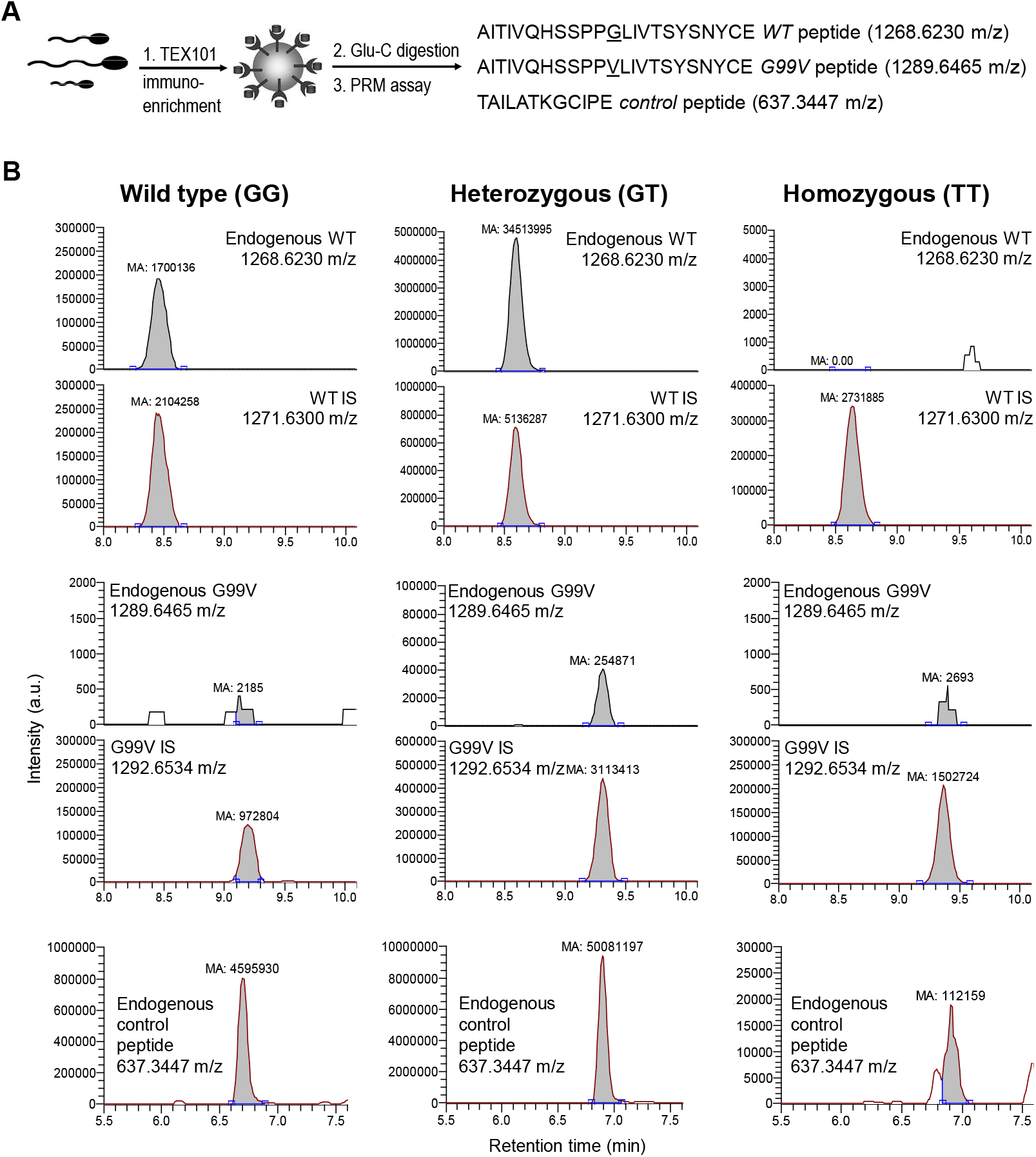
Measurement of WT and G99V variant forms of TEX101 protein by targeted mass spectrometry. (**A**) Immuno-PRM assay included immunoenrichment of endogenous TEX101 with a mouse monoclonal antibody 34ED556 coupled to sepharose beads, followed by Glu-C digestion and PRM measurements of WT and G99V peptides, as well as a control peptide representing total TEX101. (**B**) In a WT patient (GG, left panel), high levels of TEX101 were measured in seminal plasma by PRM assay without immunoenrichment. In rs35033974 heterozygote (GT, middle panel) and rs35033974^*hh*^ homozygote (TT, right panel) patients, WT, G99V variant and control peptides of TEX101 were measured by immuno-PRM assay in spermatozoa lysates. Corresponding heavy isotope-labeled peptides were used as internal standards to ensure correct identification and accurate relative quantification of peaks. Assuming theoretically equal expression of both alleles and similar ionization efficiencies of WT and G99V peptides, <3% of expected G99V variant form was found in spermatozoa of a heterozygous patient, suggesting that G99V TEX101 form may be degraded during spermatogenesis.

To explain this phenomenon, we suggested that the G99V variant form of TEX101 protein could be misfolded, aggregated and destroyed through proteasome or autophagosome degradation. A large residue of valine at position 99 could introduce substantial steric constraints, eliminate a proline-glycine beta-turn and destabilize beta-sheets in the proximity of G99V (**Supplemental Figure S2**). Interestingly, TANGO algorithm (32) for prediction of cross-beta aggregation in unfolded proteins predicted a significant impact of G99V substitution. Predicted cross-beta aggregation values changed from 3.6 for G99 to 83.8 for G99V (the average values for all residues of the mature WT TEX101 was 3.8).

### Global proteomic profiling revealed testis-specific proteins down-regulated in rs35033974^*hh*^ spermatozoa

Spermatozoa obtained from four rs35033974^*hh*^ men were considered as a TEX101 functional knockdown model. Taking into account degradation of ADAM3 proteins in spermatozoa of *Tex101* knockout mice (10), we hypothesized that TEX101-dependent human proteins would be degraded in rs35033974^*hh*^ spermatozoa and could be identified by differential proteomic profiling. Unlike immunoprecipitation approaches to identify only direct and strong physical interactions, differential profiling of the whole proteomes of WT versus rs35033974^*hh*^ spermatozoa could identify all interactions which deteriorated in the absence of TEX101 (strong, weak, direct, transient and indirect).

We thus performed a global proteomic analysis of four WT and four rs35033974^*hh*^ spermatozoa (**Figure 3A**). To achieve deep proteome coverage, peptides were subjected to the off-line fractionation by strong-cation exchange chromatography followed by the online reversed-phase liquid chromatography-mass spectrometry detection. As a result, MaxQuant analysis identified and quantified 83,984 unique peptides and 8,046 protein groups with FDR≤1.0% (Supplemental Table S4). Of 189 differentially regulated proteins (FDR≤5.0% and s0=0.4), 96 were down-regulated and 93 were up-regulated (**Figure 3B and Supplemental Table S5**). Filtering of these candidates for testis specificity using Human Protein Atlas data revealed 55 down-regulated but only 4 up-regulated proteins (potential false-positives). Thus, many more testis-specific proteins were affected by TEX101 loss. Additional filtering for cell-surface and secreted proteins using NextProt database revealed 8 down-regulated cell-surface and 9 secreted proteins, but no any up-regulated proteins (**Figure 3B**).

**Fig. 3.**
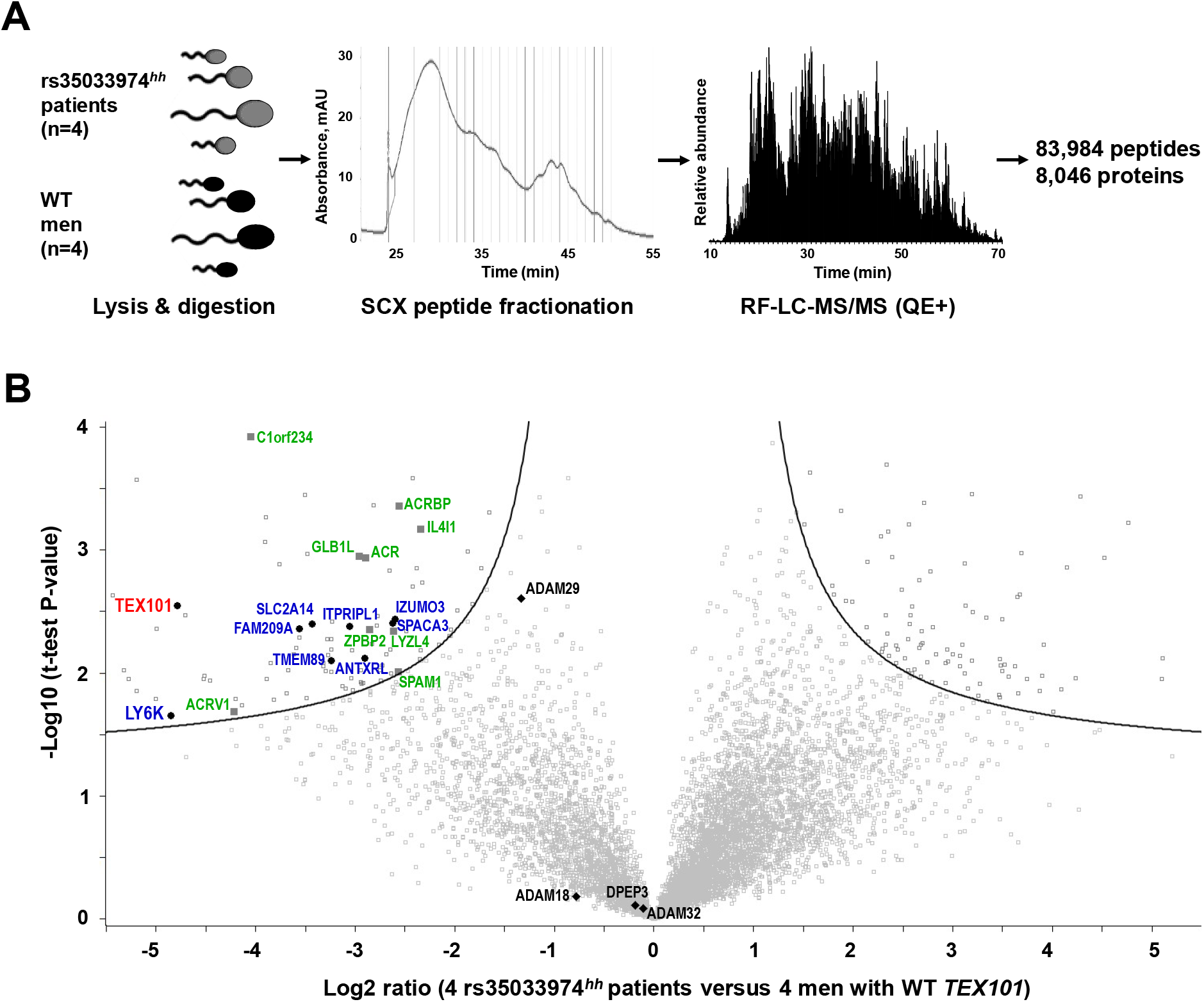
Identification of proteins down-regulated in rs35033974^*hh*^ spermatozoa, as compared to WT spermatozoa. (**A**) Spermatozoa obtained from four WT men with confirmed fertility and four rs35033974^*hh*^ patients were digested by trypsin, fractionated by strong-cation exchange chromatography, and analyzed by LC-MS/MS. (**B**) A total of 8,046 protein groups representing 8,473 unique gene names and 8,431 UniProt protein IDs were identified and quantified. Volcano plot revealed proteins down-regulated in rs35033974^*hh*^ spermatozoa with FDR ≤5% (hyperbolic curve) calculated using log2-tansformed fold change ratios, BH-adjusted *t-* test *P*-values and s0=0.4 variance correction. Differentially expressed testis-specific cell-surface and secreted proteins were plotted in blue and green, respectively. Three proteins with adhesion activity of the ADAM family (ADAM18, ADAM29, ADAM32) were identified. Levels of DPEP3, a testis-specific and TEX101-interacting protein not affected by *Tex101* knockout in mice, were not changed. No testis-specific cell-surface or secreted proteins were found significantly up-regulated.

Here, we also hypothesized that differential proteomic analysis of rs35033974^*hh*^ spermatozoa could reveal functional orthologs of mouse ADAMs 3-6 proteins destroyed in *Tex101* knockout mice (10, 12). In our spermatozoa proteome, we identified all seven human testis-specific ADAM proteins, of which three proteins with the adhesion activity (ADAM18, ADAM29 and ADAM32) could be potential orthologs of mouse ADAM 3-6 proteins (33). It was only ADAM29 protein which levels were lower in rs35033974^*hh*^ spermatozoa (**Figure 3B**). Even though ADAM29 did not pass our cut-off criteria of the global differential analysis, it was down-regulated 2.5-fold (BH-adjusted t-test *P*=0.003). We thus decided to further evaluate ADAM29 protein by SRM and Western blotting.

In this work, we identified and quantified one of the largest proteomes of human spermatozoa (8,046 protein groups representing 8,473 unique gene names and 8,431 UniProt protein IDs). Of these, 7,156 protein groups and 7,573 unique Uniprot IDs were quantified with 2 or more unique peptides. Interestingly, of 2,186 proteins currently defined as “missing” by NextProt (v2.14.0), we identified 127 proteins with 2 unique peptides (101 with previous evidence only at the transcript level, 21 inferred from homology and 5 predicted). For example, we discovered in our dataset 7 testis-elevated “missing” proteins (ANKRD60, C12orf42, LRRC63, CCDC74B, FAM47C, SPATA31A1 and TTLL8) which were identified with 2 unique peptides (≥9 amino acids) and could represent true proteins according to NextProt criteria. To conclude, our spermatozoa proteome could be used as a resource to identify and update currently “missing” testis-specific proteins.

### Evaluation of candidates by SRM and western blotting

ADAM29 and seven testis-specific cell-surface and secreted proteins involved in sperm migration (34), zona pellucida binding and penetration (35-37) and sperm-oocyte fusion (38, 39) were selected for SRM analysis. As a result of SRM measurements, 7 of 8 proteins were down-regulated in rs35033974^*hh*^ spermatozoa, while levels of DPEP3 (a testis-specific TEX101-interacting protein not affected by *Tex101* knockout in mice (11)), were not changed (**Figure 3B and Figure 4A and Table 2**).

**Table 2.** List of down-regulated testis-specific proteins in spermatozoa of rs35033974^*hh*^ homozygous men, as discovered by shotgun mass spectrometry and measured by SRM.

| UniProt accession name | Gene name | Protein | Shotgun log2-fold change | SRM log2-fold change | Protein class |
| --- | --- | --- | --- | --- | --- |
| Q9BY14 | TEX101 | Testis-expressed protein 101 | -4.8 | -4.2 | Cell-surface |
| Q17RY6 | LY6K | Lymphocyte antigen 6K | -4.9 | -3.8 | Cell-surface |
| P10323 | ACR | Acrosin | -2.9 | -3.6 | Secreted |
| Q6X784 | ZPBP2 | Zona pellucida-binding protein 2 | -2.9 | -1.2 | Secreted |
| Q8IXA5 | SPACA3 | Sperm acrosome membrane-associated protein 3 | -2.6 | -2.1 | Cell-surface |
| Q5VZ72 | IZUMO3 | Izumo sperm-egg fusion protein 3 | -2.6 | -1.2 | Cell-surface |
| P38567 | SPAM1 | Hyaluronidase PH-20 | -2.6 | -3.3 | Secreted |
| Q8NEB7 | ACRBP | Acrosin-binding protein | -2.6 | -2.9 | Secreted |
| Q9UKF5 | ADAM29 | Disintegrin and metalloproteinase domain-containing protein 29 | -1.3 | -1.2 | Cell-surface |
| Q9H4B8 | DPEP3 | Dipeptidase 3 | -0.2 | 0.4 | Cell-surface |

**Figure 4.**
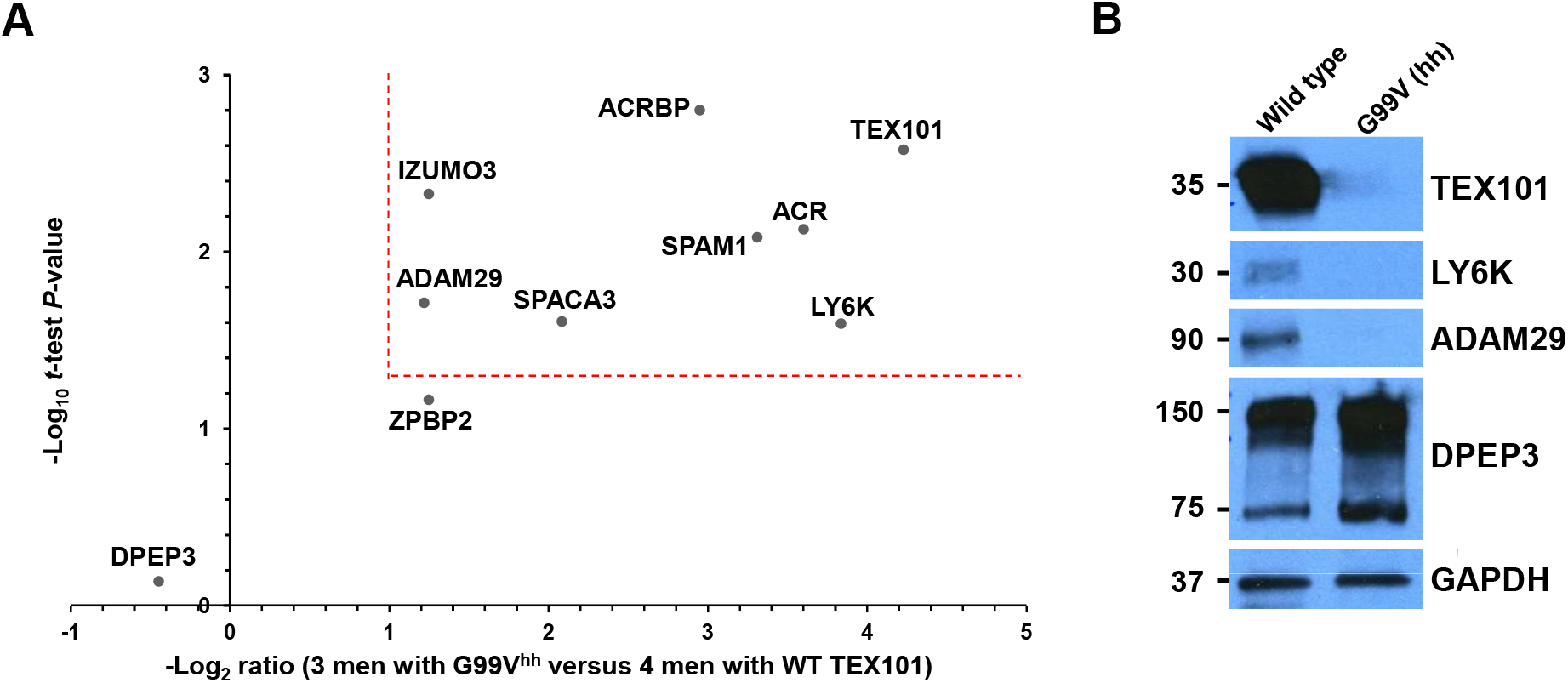
Verification of 8 testis-specific cell-surface and secreted proteins in four WT and three rs35033974^*hh*^ men by SRM and western blotting. (**A**) A multiplex SRM assay facilitated accurate relative quantification of 8 candidate proteins and confirmed significant down-regulation (>2-fold change, P-value <0.05, dotted lines) of 7 proteins. (**B**) Western blot analysis of spermatozoa obtained from one WT and one rs35033974^*hh*^ men confirmed reduced levels of TEX101, LY6K and ADAM29 proteins, while levels of a testis-specific protein DPEP3, present in spermatozoa as both a monomer and a homodimer, were not affected. GAPDH was used as a loading control for total protein.

Interestingly, the top down-regulated protein in rs35033974^*hh*^ spermatozoa was LY6K (14-fold change; *t*-test *P*-value=0.03). LY6K (Lymphocyte antigen 6K) is a testis-specific GPI-anchored protein localized at the cell surface of testicular germ cells, with a similar expression pattern to TEX101 based on immunohistochemistry data available at the Human Protein Atlas. It was previously demonstrated in mice that LY6K physically interacted with TEX101, but disappeared from the cell surface in the absence of TEX101 protein in *Tex101* knockout mice (11, 34). Finally, we evaluated LY6K, ADAM29 and TEX101 proteins by western blotting. Levels of these proteins were undetectable in rs35033974^*hh*^ spermatozoa, while expression of DPEP3 monomers (75 kDa) and dimmers (150 kDa) was found at normal levels (**Figure 4B**).

## DISCUSSION

It is estimated that 30% of infertile men are diagnosed with unexplained infertility, often with normal semen parameters of sperm count and motility (40). The routine testing for male infertility is currently limited to analysis of sperm concentration, motility and morphology (41), as well analysis of few genetic abnormalities including Y chromosome deletions, CFTR mutations and sperm DNA fragmentation for specific indications (42). Many more genetic abnormalities associated with male infertility have been identified (43-47), however, improved diagnostics of male infertility remains an unmet need. Elucidation of functional roles of testis-specific proteins may not only advance biology of human reproduction, but also provide novel diagnostic markers, particularly for male infertility subtypes which cannot currently be explained with routine diagnostics.

To identify genes essential for male fertility, approximately 500 knockout mouse models have been investigated to date. Some models resulted in male infertility with severe defects in spermatogenesis, spermiogenesis and sperm maturation, or infertility with no apparent defects in motility, morphology and sperm count (*Ace*, *Adam2 and Adam3*) (48, 49). Other models revealed sub-fertile phenotypes with reduced acrosome reaction and capacitation (*Acr*, *Pcsk4*), decreased motility (*Smcp*), delayed cumulus-oocyte complex dispersion (*Spam1*), weak zona pellucida binding (*Press21*) and reduced sperm-oocyte fusion (*Crisp1*) (50). Surprisingly, knockout studies in mice also revealed numerous sperm cell surface proteins not essential for fertilization *in vivo*, thus suggesting compensatory mechanisms and multi-factorial nature of infertility (50).

Recently, the Human Protein Atlas profiled testis-specific genes and proteins, and identified 1,079 genes with more than five-fold higher mRNA levels in testis as compared to all other human tissues (1). Due to their exclusive expression in testis, these proteins may be essential for spermatogenesis, remodeling of sperm surface proteome, sperm transit and sperm-oocyte fusion (51). However, lack of cell lines expressing testis-specific proteins was the major bottleneck to study the molecular function of human testis-specific proteins *in vitro*. Likewise, primary human germ cells isolated from orchiectomy samples could not be maintained in long-term cultures and studied with gene knockout or knockdown approaches.

Identification of natural ‘human knockouts’ with homozygous loss-of-function mutations provided an alternative approach to study functional and pathological roles of human proteins (52). The most valuable mutations for such studies included protein truncating variants due to stop gain or frameshift mutations, or single amino acid variants leading to the loss of activity or to protein misfolding followed by proteasomal degradation. In this work, we hypothesized that spermatozoa of men with natural knockouts or functional knockdowns of testis-specific genes could emerge as valuable models to study functional and pathological roles of human testis-specific proteins.

Recent population-scale studies of genetic variation (25, 53, 54) discovered numerous protein truncating variants or single amino acid variants in humans, and provided their accurate frequencies in different ethnic groups (55). For example, the Genome Aggregation Database (www.gnomad.broadinstitute.org) contained the whole-exome and whole-genome sequencing data for 138,632 individuals with loss-of-function mutations and predicted deleterious impact of missense variants with Polyphen an SIFT algorithms (25). Variants present with the minor allele frequencies ≥0.3% and 5.5% could be of particular interest since such frequencies ensure identification of at least 3 heterozygous or homozygous individuals, respectively, per 1,000 individuals in the general population. It is expected that frequencies of pathogenic variants will be further enriched in patient cohorts, such as men referred to fertility clinics. Thus, the impact of truncating or single amino acid variants on the expression or activity of corresponding testis-specific proteins could be investigated experimentally in clinical samples, such as spermatozoa or testicular tissues. Since immunoassays are not available for the majority of testis-specific proteins and also typically cannot detect single amino acid substitutions, mass spectrometry-based approaches remain the only tools to measure the expression of variant proteins (56, 57).

In this study, we focused on a testis-specific protein TEX101 which we previously identified and validated as a seminal plasma biomarker for the differential diagnosis of azoospermia and male infertility (58-60). TEX101 was previously shown to be essential for the production of fertilization-competent spermatozoa in mice (10, 12, 61-63). Targeted knockouts of *Tex101* gene resulted in absolute male sterility, with no other phenotypic abnormalities and normal sperm concentration and morphology (10, 11). The presence of TEX101 protein was shown to be crucial for the trafficking, maturation or localization of four proteins of the ADAM family (ADAM3, ADAM4, ADAM5 and ADAM6) essential for fertilization in mice (10-12). However, mouse data could not be translated into human studies since human ADAM3, ADAM5 and ADAM6 were non-coding pseudogenes, while ADAM4 was not present in the human genome (13).

Here, we searched population-scale genomic databases for the loss-of-function protein truncating and single amino acid variants of human *TEX101* gene, identified rs35033974 variant and discovered substantially lower levels of the G99V variant form of TEX101 protein in spermatozoa of heterozygous and homozygous men. We then hypothesized that rs35033974^*hh*^ spermatozoa could be used as a knockdown model to study the molecular function of human TEX101, discover its functional interactome and identify orthologs of mouse ADAMs 3-6. Our proteomic approach was designed to identify the TEX101 functional interactome, such as proteins co-degraded together with G99V TEX101 in rs35033974^*hh*^ spermatozoa. We hypothesized that some of these interactions could be weak or transient and thus missed by typical co-immunoprecipitation approaches.

As a result, we identified and verified 8 cell-surface and 9 secreted testis-specific proteins which were significantly down-regulated in rs35033974^*hh*^ spermatozoa. It should be emphasized that identification of LY6K protein as a top candidate suggested the robustness of our experimental protocol. Indeed, previous studies in mice revealed that LY6K physically interacted with TEX101 and disappeared from the spermatozoa surface in *Tex101* knockout mice (62).

Similar to TEX101, LY6K is GPI-anchored cell-surface protein expressed by testicular germ cells and is partially shed into seminal plasma during sperm maturation (34). Interaction of TEX101 with LY6K in mice was shown to be crucial for proper trafficking and post-translational processing of LY6K (11). *Tex101*^-/-^ or *Ly6k*^-/-^ mice were infertile due to compromised migration of sperm in the oviduct (10, 11, 34). TEX101-LY6K complex facilitated proper processing of ADAM3 protein. Interestingly, levels of TEX101 and LY6K proteins on the surface of spermatozoa, but not levels of intracellular mRNA transcripts, were mutually dependent. Thus, LY6K protein quickly degraded in *Tex101*^-/-^ mice, and vice versa (11). Interestingly, another cell-surface GPI-anchored protein, a testis-specific dipeptidase DPEP3, also formed a physical complex with TEX101 (63), but DPEP3 levels were not affected in *Tex101*^-/-^ mice (11). Thus, our data on human TEX101, LY6K and DPEP3 in WT and rs35033974^*hh*^ spermatozoa (**Figure 4**) confirmed previous observations in mice.

Global proteomic profiling of spermatozoa from four WT and four rs35033974^*hh*^ men identified eight testis-specific ADAM proteins with adhesion (ADAM2, ADAM18, ADAM29, ADAM32) and metalloprotease (ADAM20, ADAM21, ADAM28, ADAM30) activities (33). Interestingly, it was only ADAM29 which levels decreased in rs35033974^*hh*^ men, as discovered by global proteomic profiling (2.5-fold, *P*=0.003) and verified by SRM and western blotting. Since molecular function of human ADAM29 protein has never been previously reported, we suggest that ADAM29 protein should be further investigated as one of the potential functional orthologs of mouse ADAM 3-6 proteins.

We also suggested here that very low levels of G99V variant form in heterozygous and homozygous spermatozoa (~3% of predicted) could be the result of misfolding and subsequent proteasomal degradation (64-66). Similar impact was previously observed for misfolded CFTR (67) and some GPI-anchored proteins (68). Introduction of a large hydrophobic valine residue at position 99 within the PP<u>G</u>L beta-turn could result in substantial steric constraints, destabilization of proximal beta-sheets and subsequent protein aggregation (69, 70). Significant impact of G99V substitution on beta-aggregation was in agreement with TANGO calculations (32). The predicted TANGO impact was the most deleterious for substitutions of G99 with hydrophobic residues such as valine, isoleucine and phenylalanine. Glycine at position 99 was also found highly conserved in mammals (**Supplemental Figure S1**).

In this work, we identified and quantified one of the largest proteomes of human spermatozoa (8,046 protein groups representing 8,473 unique gene names and 8,431 UniProt protein IDs). At least 7 testis-elevated proteins (ANKRD60, C12orf42, LRRC63, CCDC74B, FAM47C, SPATA31A1 and TTLL8) identified in our work were previously considered “missing proteins” in NextProt, UniProt and Human Protein Atlas databases. Our deep proteome of spermatozoa could be used as a resource to update currently “missing” testis-expressed proteins.

Finally, we speculated on possible deleterious impact of rs35033974 allele on male fertility. Even though TEX101 protein levels were found significantly lower in rs35033974 heterozygous men, 70% of heterozygous men still had TEX101 levels in seminal plasma above 65 ng/mL, a cut-off level of male infertility which we previously established (8). Due to its relatively high allele frequency in the general population (12.4% in Europeans), rs35033974 is unlikely a severe pathogenic factor leading to male sterility. In addition, if rs35033974 would be a recessive pathogenic variant with a high penetrance, the number of homozygous men in population would be substantially lower than that calculated using the Hardy-Weinberg equilibrium. We found no substantial deviation from the equilibrium for the European population in gnomAD (*N*=63,332; 1,011 homozygotes identified versus 982 calculated). According to the 1000 Genomes Project, the rs35033974 variant was present with the similar allele frequencies in males and females. Interestingly, 1000 Genomes Project data (26) included 9 homozygous men, of which 5 had biological children (see examples in **Supplemental Figure S3**). Since TEX101 levels were found significantly lower in heterozygous and homozygous men, rs35033974 status may be considered in the use of TEX101 protein as a male infertility biomarker. In addition, patients with different ethnic backgrounds may have different cut-off values for TEX101, with lower cut-offs for European men, of which 21.8% are heterozygous and 1.6% are homozygous.

There may be several possibilities to explain the discrepancy between molecular and clinical data on TEX101 protein: (i) unlike mouse TEX101, human TEX101 protein and TEX101-LY6K interaction may not be essential for sperm maturation, ADAM processing and fertilization; (ii) low levels of TEX101 protein in rs35033974^*hh*^ patients may be compensated by alternative cell-surface chaperons; (iii) rs35033974 is deleterious for protein structure, however, unlike mouse *Tex101*, human *TEX101* could be another non-essential gene (71, 72). Future studies should investigate if this highly-frequent variant (1.6% homozygous genotype frequency in European population) predisposes males to infertility and becomes pathogenic in combination with other factors, for example, lowered sperm concentration in semen.

It should be noted that our study had the following limitations: (i) even though label-free quantification using MaxQuant algorithm is recognized as an accurate proteome-wide quantification approach (16), its variability may still be relatively high, so all candidates should be verified by orthogonal assays, such as SRM or western blotting; (ii) global proteomic quantification of a very large number of proteins (8,046) and FDR-based cut-offs could result in many false-positive (for example, intracellular non-testis-specific proteins) and false-negative candidates (ADAM29 could be such a false-negative candidate); (iii) while the results of our global proteomic quantification were confirmed by SRM assays in three rs35033974^*hh*^ and four WT patients, and by western blotting in single rs35033974^*hh*^ and WT patients, we could not recruit additional rs35033974^*hh*^ patients for further validation of our data; (iv) fertility status of our four rs35033974^*hh*^ patients is not known. One rs35033974^*hh*^ patient proceeded to infertility treatment by intrauterine insemination, but the treatment of that couple was not successful (female factor, however, was not excluded).

To conclude, we presented the first human study to investigate the possible functional role of TEX101 protein as a cell-surface chaperone, and identified degradation of LY6K, as well as additional 7 cell-surface testis-specific proteins, in the absence of TEX101. Spermatozoa of rs35033974^*hh*^ men may be used as a unique model to elucidate further details on the role of human TEX101 which will advance male reproduction biology. Since TEX101 seminal plasma levels were found significantly lower in heterozygous than in wild-type men, rs35033974 status could be considered in TEX101 diagnostics. Further studies on rs35033974 and its association with male infertility diagnosis or prognosis will be required. Our work may serve as a concept for future studies on functional effects of natural knockouts or knockdowns in humans and could lead to development of novel male infertility diagnostics (73, 74) and discovery of targets for male contraceptives (75). Presented approach may also facilitate verification of the essential and non-essential testis-specific genes and proteins, which will advance biology of human reproduction.

## Acknowledgements

We thank Ihor Batruch for assistance with mass spectrometry, and Susan Lau for coordinating collection and storage of clinical samples.

## Author contributions

A.P.D., E.P.D. and K.J. designed the research project; C.S. performed the experiments; C.S. and A.P.D. analyzed the data. D.K. produced anti-TEX101 mouse monoclonal antibodies, and K.J. provided clinical samples and clinical expertise. C.S. and A.P.D. wrote the manuscript, and all authors contributed to the revision of the manuscript.

## Supplemental materials

This article contains supplemental materials.

